# Glucosylceramide depletion disrupts endolysosomal function in GBA-linked Parkinson’s fibroblasts?

**DOI:** 10.1101/2023.11.30.569349

**Authors:** M Bhardwaj, Z Kula, Y Weng, D te Vruchte, C Breda, D.J. Sillence

**Affiliations:** De Montfort University; Oxford University

## Abstract

In Gaucher and Niemann-Pick C diseases, the glucosylceramide (GlcCer) depletion hypothesis states that depletion of non-lysosomal sphingolipid pools can lead to dysfunction in the secretory and lysosomal system. The hypothesis suggests: 1) lysosomal dysfunction can be separated from lysosomal storage, 2) Lysosomal/secretory dysfunction/vATPase activity is corrected by increasing non-lysosomal GlcCer pools, and 3) Changes in higher glycosphingolipid synthesis due to changes in Golgi pH and/or GlcCer non-vesicular transport. Evidence for this mechanism includes 1) Successful treatment of cells and animals by imino sugar inhibition of the non-lysosomal neutral pH GlcCer hydrolase GBA2, 2) Increasing ER/cytosol GlcCer increases in vATPase regulatory V0a1 subunit expression.

Heterozygous mutations in GBA1, a lysosomal glucocerebrosidase (GCase), cause GCase misfolding and mislocalisation in the ER/cytoplasm which is linked to Parkinson’s disease (GBA-PD). Unexpectedly, similar to previous results in storing fibroblasts, N370S and L444P fibroblasts revealed increased endolysosomal pH and size despite the absence of glucolipid storage. Induction of storage by reducing residual lysosomal GCase activity in the N370S/L444P fibroblasts by the addition conduritol B-epoxide had no further effect on lysosomal function. In contrast, the addition of a soluble GlcCer analogue (adaGlcCer) reverses increased endolysosomal pH and volume in N370S mutant fibroblasts. The results are consistent with ER/cytosolic glucolipid depletion in GBA-PD fibroblasts. We discuss the potential for toxic/ectopic GBA1 hydrolysis and disrupted vATPase activity may lead to defective dopamine packaging and synaptic vesicle endocytosis as a new hypothesis in GBA-PD.

## Introduction

In Parkinson’s disease, multiple risk factors such as synuclein, LRRK2, PARKIN for have been suggested to be unified in a general defect in synaptic vesicle endocytosis and vATPase-mediated dopamine packaging (1,2). Genetic mutations in the GBA1 gene increase the risk of developing Parkinson’s disease (PD). However, it remains to be established how GCase misfolding leads to dopaminergic dysfunction. GlcCer is degraded by glucosylceramidase (GCase) to ceramide and glucose by acid GCase hydrolysis is in the lysosomal lumen. However, glucosylceramide (GlcCer) is widely distributed throughout the ER and plasma membrane where it is degraded (3,4) by a non-lysosomal neutral GBA2 (5) GlcCer has vital roles. In melanocytes, Gaucher macrophages and NPC fibroblasts, ER/cytosolic-facing GlcCer modulates vATPase in the Golgi and lysosomes (6–8) which is necessary for correct tyrosinase and lipid sorting (9–11). In the group of PD patients where *GBA* is affected, 73% carry mono-allelic L444P and N370S mutations (12). Clearly, homozygous GBA1 mutations are linked to reduced lysosomal activity and cause lysosomal storage, Gaucher disease (GD). In contrast, no GlcCer storage has been found in GBA-PD (13,14) and some GBA-PD mutations are not associated with Gaucher disease (E236K) suggesting distinct mechanisms (15). A clinical trial was performed for substrate reduction therapy of GBA-PD patients using venglustat to inhibit GlcCer synthesis and reduce lysosomal storage. However, this trial was unsuccessful as treatment was associated with the acceleration of PD symptoms (16,17). This result suggests that GlcCer storage does not cause PD symptoms and is instead necessary for appropriate dopaminergic function.

In fibroblasts from PD patients, lysosomal pH is increased (18–20). Previous studies have suggested that cytosol-facing endomembrane pools of GlcCer are linked to the regulation of lysosomal pH and vATPase activity or expression (6–9,21). Since cytosolic GlcCer is linked to lysosomal acidification, we sought possible toxic gain of function of misfolded GCase in non-storing skin fibroblasts, due to the unregulated hydrolysis of non-lysosomal GlcCer. It has been reported that GBA2 inhibition leads to positive effects in Gaucher and related diseases (22–24). This implies that non-lysosomal cytosol-facing/ER GlcCer is needed for correct cell function. Preliminary data agreed with previously published reports of slightly decreased GBA2 expression in Gaucher fibroblasts (25–27). In the present report and in agreement with previous results N370S/L444P fibroblasts do not generally store GlcCer or GlcSph (11,28). In contrast, some L444P fibroblasts did show significant changes in a-series gangliosides possibly due to changes in non-vesicular transport of GlcCer (13,29). Incubation with conduritol B-epoxide (CBE), a pH independent GCase inhibitor, increased storage but did not lead to further increases in lysosomal pH or size in L444P/N370S fibroblasts. In contrast, treatment with a soluble GlcCer analogue, adamantyl GlcCer (adaGlcCer) (30) resulted in reversal of increased endolysosomal pH and size in N370S/V394L fibroblasts. In contrast, GCase activation with the non-inhibitory chaperone adamantlylGalCer did not change endolysosomal pH or size. Our results support the hypothesis that mono-allelic misfolded/mislocalised residual GCase activity can lead to GlcCer depletion. A hypothetical model is discussed where toxic GCase mislocalisation and ectopic hydrolysis of inappropriate ER/cytoplasmic GlcCer pools leads to disrupted vATPase resulting in deficient dopamine packaging during synaptic vesicle recycling in GBA-PD.

## Results

### GlcCer-depletion as an important mechanism for Gaucher and Niemann-Pick C (NPC1) disease

We have shown that endolysosomal defects in Niemann-Pick C fibroblasts are corrected by inhibition of cytosolic GBA2 (6). For instance, U18666A (a putative inhibitor of NPC1) increases the size of endolysosomes. Correcting deficient cytosolic-facing GlcCer by inhibition of GBA2 with 10nM AMP-DNJ reverses the defect (Fig. 1A).

**Figure 1.**
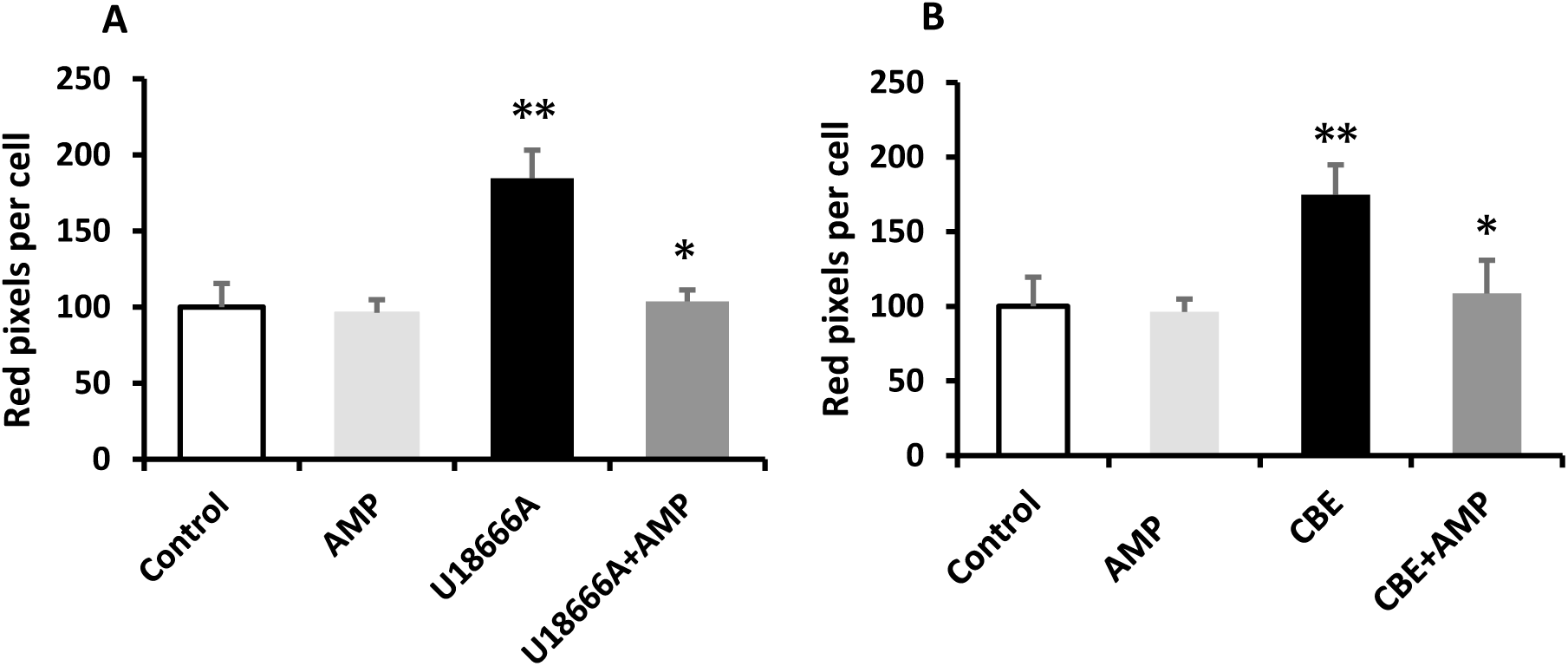
ER/Cytosolic GBA2 modulates endolysosomal volume in macrophages. RAW macrophages were labelled with 150nM LysoTracker red DND-99 in 1% RPMI serum media for 15 minutes at 37^0^C. (A) Quantitation of punctate size in RAW cells in the presence of 300nM U18666A incubated with 10nM AMP-DNJ overnight (cytosolic GBA2 inhibitor) (B) Measurement of punctate size in RAW macrophages in the presence of 25uM CBE and incubated with 10nM AMP-DNJ overnight. (* *p* < .05, ** *p* < .001 *n* = 3, *t*-test). Data were normalised to the control. Results are presented as mean ± SD.

We initially expanded these observations in a cell culture model of Gaucher disease (31–33). 25µM CBE treatment increases endolysosomal size in mouse macrophages (control 100 to 170 ± 0.1 CBE treated: P=0.01), consistent with increased storage observed in Gaucher macrophages (11,34). The inhibition of GBA2 by 10nM AMP-DNJ significantly reverses the endolysosomal size (Fig. 1B).

### A soluble GlcCer mimetic corrects endolysosomal defects in N370S GBA1 fibroblasts

Initial results in Gaucher fibroblasts suggested that endolysosomal pH is increased in a variety of N370S and L444P fibroblasts. This was unexpected as Gaucher fibroblasts are reported not to store since residual enzyme activity is considered high enough to degrade lysosomal GlcCer (11,28,35). In order to investigate whether these increases in pH were due to GlcCer-depletion in a similar fashion to storing macrophages, a soluble short chain GlcCer derivative adamantyl-GlcCer (adaGlcCer) was used (30). The endolysosomal pH and size was reversed in N370S/V394L fibroblasts treated with 40 µM adaGlcCer overnight (Fig 2). However, no changes in pH or size were observed in the wtGCase control. As a further control the galactose analogue (a non-inhibitory GCase chaperone) adaGalCer (30) had no effect (Fig. 2).

**Figure 2.**
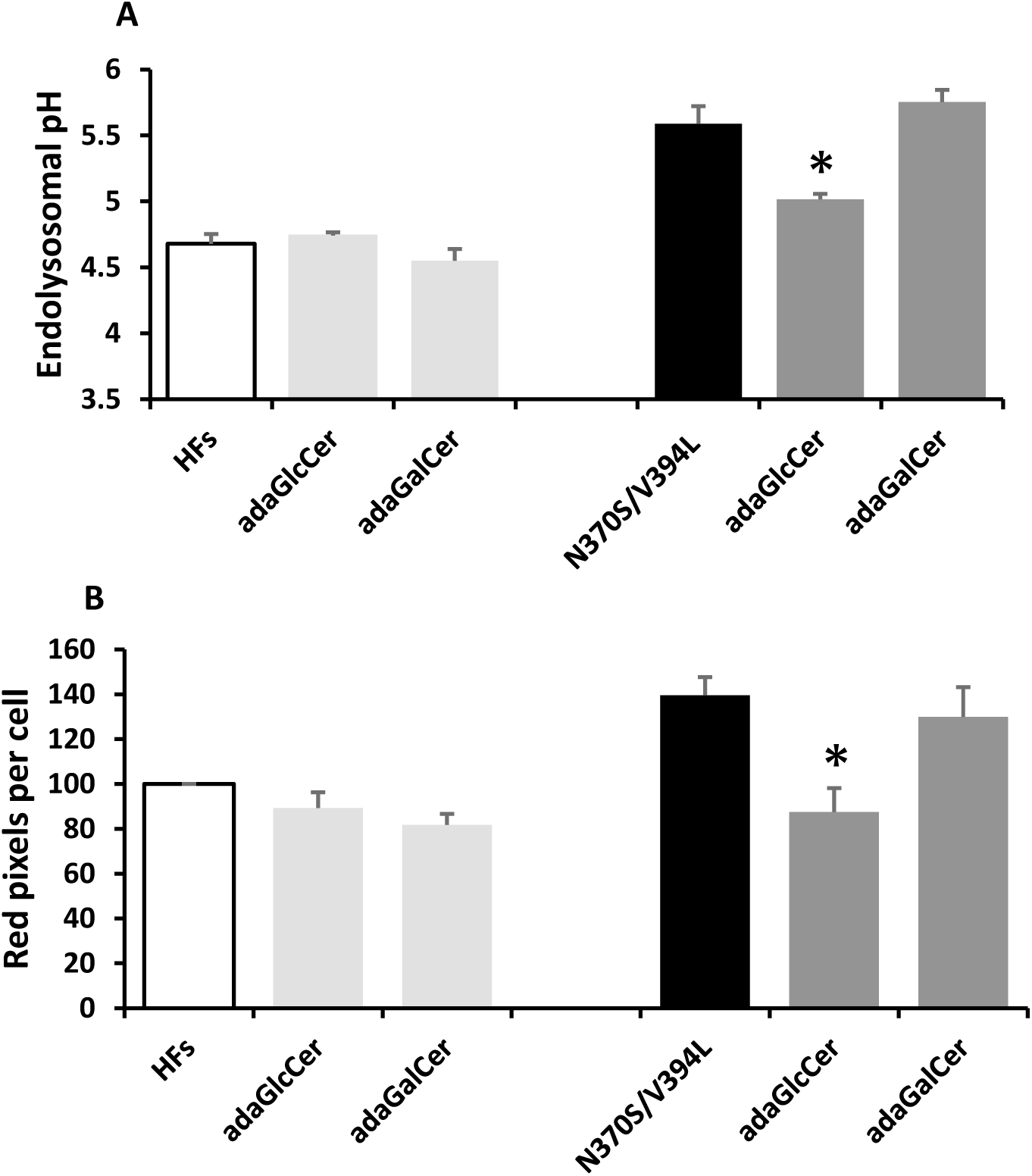
Adamantyl-GlcCer decreases endolysosomal pH and size in non-storing N370S fibroblasts. (A) Overnight treatment with 40 µM adaGlcCer significantly reduces endolysosomal pH (lysosensor Y/B) in N370S patient fibroblasts, n=2-5 (B) Quantitation of endolysosomal size (Lysotracker red) showed significant reversal in N370S patient fibroblasts with 40µM adaGlcCer n=3-5. Results are mean of ±SD. Significance measured by t-test.

### GBA1 mutant fibroblasts do not store glucolipids

The most obvious explanation for the observed increases in endolysosomal size and pH was that L444P/N370S fibroblasts were storing GlcCer. In order to investigate this further, we measured the levels of GlcCer (3) and glucosylsphingosine (GlcSph) (36) in a variety of mutant fibroblasts. GBA1KO fibroblasts have 33 times greater GlcCer and 2-fold GlcSph compared to controls (Fig. 3A). In contrast in a L444P and N370S fibroblasts (residual GCase activity between 3% and 25%) (37), no significant increase in GlcCer/GlcSph levels were detected, except the most severe homozygote (L444P/L444P). The addition of a GBA2 inhibitor (AMP-DNJ) to GBA1KO fibroblast did not increase glucolipid levels any further (up to 24 hour) suggesting that the cytosol/ER GlcCer is a minor pool. This contrasts with mouse brain reports in which GlcCer levels is increased >20 fold upon imino sugar treatment (38) likely due to much higher levels of GBA2 expression in the brain.

**Figure 3.**
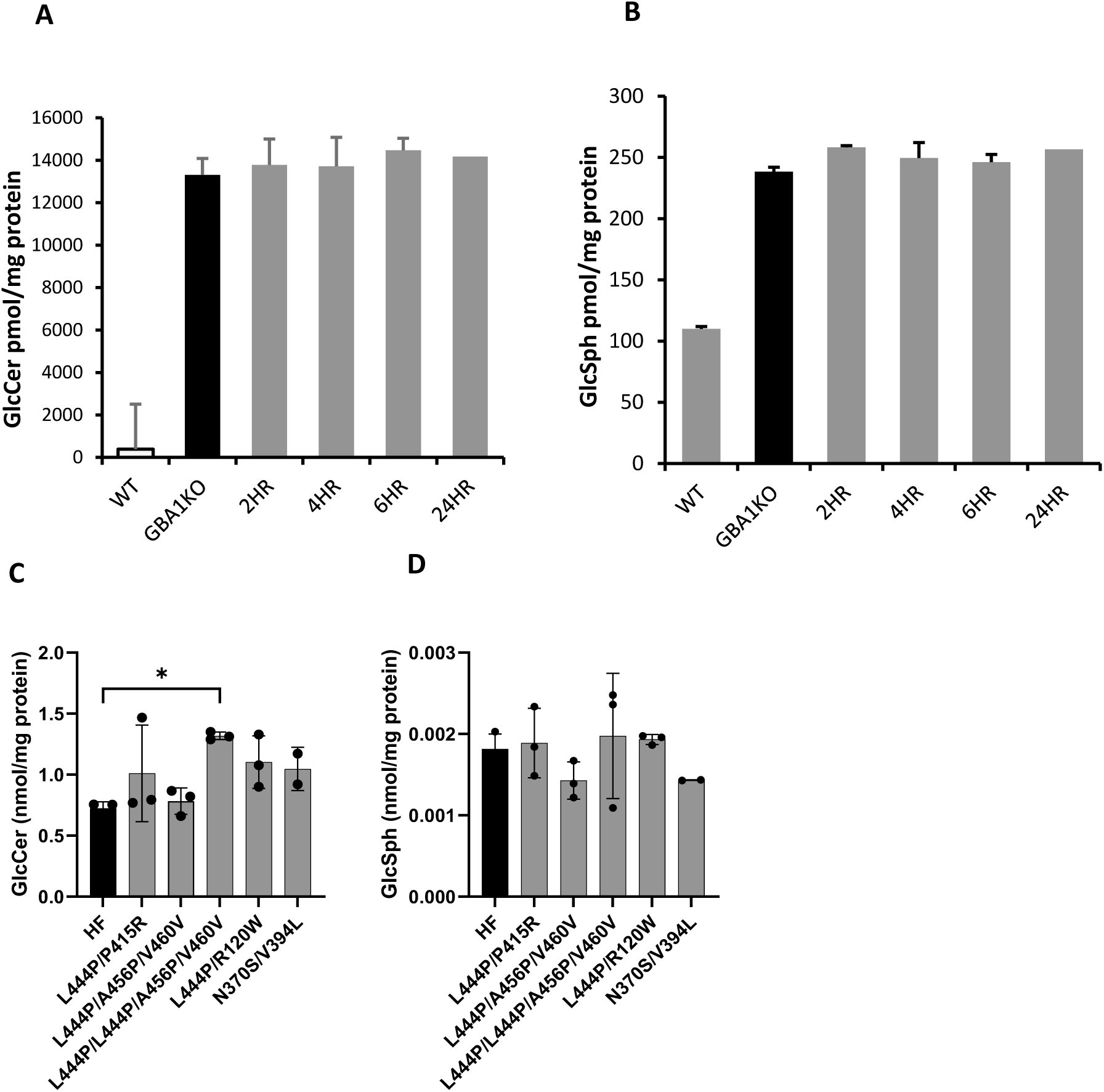
Measurement of N370S/L444P fibroblasts sphingolipids levels. Quantitative lipid analysis by mass spectrometry. Total cell lysate was used to determine lipid levels (A, B) HAP GBA1 knockout were treated with 5nM AMP DNJ for 2, 4, 6 and 24 hours. GlcCer and GlcSph levels compared with time-dependent inhibition of GBA2. (C, D) Glycosphingolipids comparison between healthy control and GBA1 mutants. Data is represented as mean ±SD, n=3. Significance measured by t-test and ANOVA. *\*p ≤ 0.05, **p < 0.01, ***p < 0.001 and ****p < 0.001*. A lipid analysis was conducted by Danielle Taylor-Te Vruchte and Yuzhe Weng (University of Oxford).

Changes in higher glycolipid levels have also been observed in GBA-PD (13,36). Despite little change in glucolipids higher a-series glycosphingolipids (GSLs) were found to be increased in several L444P fibroblast cell lines. Changes in higher a series glycosphingolipids in PD have been suggested to be consistent with altered glycolipid synthesis rather than breakdown (13,36). It should be noted that higher glycosphingolipid synthesis also relies on non-vesicular transport of cytosolic GlcCer which may be cell-type dependent (29).

### Fibroblasts with GBA-PD mutations are resistant to further increases in GlcCer

Given the very low or complete lack of storage that occurs in the N370S/L444P fibroblasts, storage was induced with the addition of CBE which would be expected to inhibit any residual lysosomal enzyme. Overnight treatment with 25 µM CBE did induce storage with 10 fold increases in GlcSph (Table S1). Despite the induction of storage, no additional changes in endolysosomal pH were observed in N370S/L444P patient fibroblasts with 25µM CBE treatment for 24 hours (Figure 4B), despite a CBE-induced increase in pH in wt fibroblasts. Additionally, CBE treatment did not lead to any increase/decrease in lysosomal volume in N370S/L444P fibroblasts (Fig. 4B). The results are therefore consistent with the hypothesis that increased endolysosomal pH and volume in the L444P/N370S fibroblasts are due to altered non-lysosomal GlcCer.

**Figure 4.**
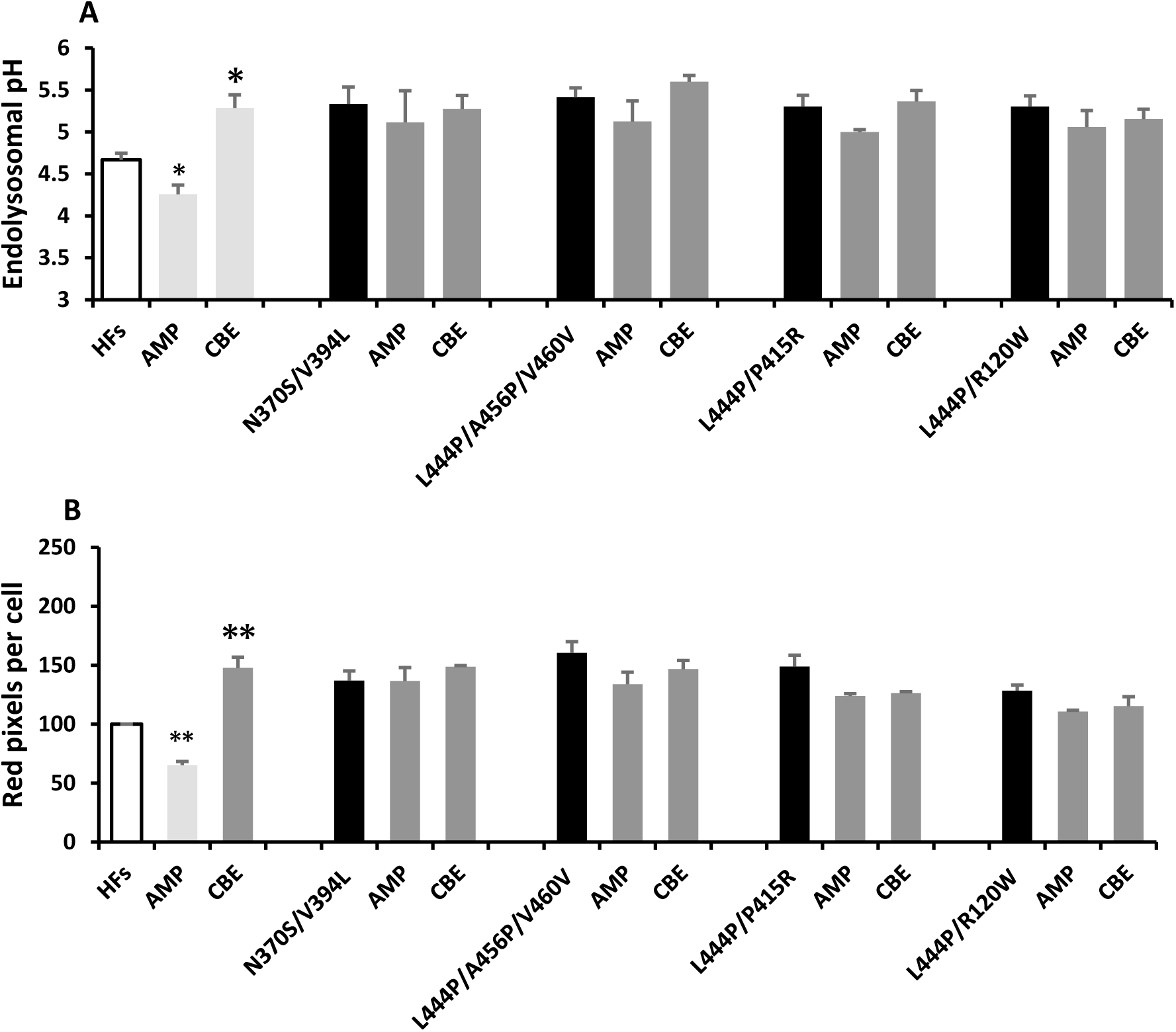
GBA-PD fibroblasts are resistant to further increases in GlcCer. (A) No reduction and increase in endolysosomal pH were observed in various Gaucher patient fibroblasts with 10nM AMP-DNJ (leads to significant increase in sphingosine and ceramide) and 25uM CBE overnight treatment (10 fold increase in GlcSph), respectively. n=2-12, (B) GD/PD fibroblasts showed increased endolysosomal volume/size that was not changed on GBA2 inhibition with 10nM AMP-DNJ or 25 uM CBE, n=2 All data are represented as mean ±S.D. Significance measured by t-test and ANOVA.

## Discussion

In the present study, we find evidence for the GlcCer-depletion hypothesis for GBA-PD exploiting a panel of Gaucher fibroblasts carrying PD-associated mutations. In agreement with previous reports no accumulation of GlcCer or GlcSph were observed apart from the most severe L444P/L444P mutation. Despite no storage, similar to other GlcCer-depleted cells using inhibitors of GlcCer synthesis (6,7,11) impaired endolysosomal pH and size are observed with the GBA-PD mutations similar to previous studies (19,20). Consistent with GlcCer-depletion, endolysosomal defects were corrected by the addition of adamantyl-GlcCer but not the GalCer analogue, reminiscent of lactosylceramide trafficking in GlcCer-depleted macrophages (11). Non-lysosomal GlcCer depletion suggests a new mechanism for GBA1-related changes in acidification within the secretory and lysosomal system in a similar fashion to that observed in other cells (6,8). The importance of defective vesicular acidification is still an open question but includes the following: 1) increased pH may decrease dopamine transport and packaging during synaptic vesicle endocytosis which has been proposed as a central mechanism in PD (1,2), 2) mislocalised tyrosinase transport and neuromelanin synthesis is GlcCer-dependent (8,9), although this may be less likely (39). It should also be noted that apart from regulation of vATPase ER GlcCer depletion has also been linked to dysfunction in ERAD/retrotranslocation through the disruption of ER GlcCer rafts (3). Such an effect may well be expected to exacerbate ER stress that’s one of features in GBA-PD (20,40).

It should be noted that so far, this hypothesis remains unproved and is presented to further discussion in the field. To date, it has not been possible to measure decreases in either GlcCer or GlcSph in whole PD cells. We think it likely that the relevant non-lysosomal GlcCer pool is a minor signalling pool. We have also not been able to measure any increase in either GlcCer or GlcSph whole cell levels on inhibition of GBA2 in wt fibroblasts (although sphingosine and ceramide do increase). Even with low GBA2 expression, inhibition of this enzyme leads to significant changes in pH and vATPase expression (6). In fibroblasts, inhibition of GBA2 also seems to regulate levels of membrane contact sites between lysosomes and ER, consistent with MCS being the location of a minor lipid signalling pool (Sillence/Eden unpublished).

The mechanism of action of soluble adamantyl GlcCer in the presented experiments is an open question and could be simply mimicking GlcCer in the cytoplasm. However, we favour adamantylGlcCer is acting through inhibition of GBA1 in the ER/cytoplasm at neutral pH (30) for the following reasons: 1) adaGlcCer treatment had no effect on L444P fibroblasts (Fig. S2) and did not inhibit GBA1 activity in fibroblast homogenates (not shown), 2) adaGlcCer did not have an effect on wt fibroblasts where GBA1 was not expected to be mislocalised to the ER/cytosol, 3) pH-independent inhibition by Conduritol B-epoxide (CBE) has no effect due to equal inhibition in the lysosome and the ER which balance each other out. 4) Inhibition of ER/cytosol GBA2 has no effect which is consistent with continued GlcCer hydrolysis by ER GCase activity in the L444P/N370S fibroblasts 5) A role for both GBA1 and it’s pseudogene in the ER has recently been proposed (41).

GlcCer depletion due to mislocalised GCase activity in the ER/cytoplasm is also favoured for the following reasons 1) GlcCer is present throughout the cell including the ER (3,4) and regulates vATPase (7,8,21,24). 2) Excess hydrolysis and inappropriate ectopic hydrolysis provides a mechanism for lysosomal dysfunction in the absence of storage. 3) N370S GCase is potentially active in the ER/cytosol due to an increased pH optimum of the misfolded protein, especially in the absence of SAP activators (42–44). It could also be noted that unsaturated negatively charged ER phospholipids may support mislocalised GCase activity (45,46).

In contrast to the above in the storing L444P/L444P fibroblasts severe homozygous mutant of GBA1 showed significant increases in endolysosomal pH (5.4±0.06 *(p=0.0001)* which are reversed upon GBA2 inhibition with AMP-DNJ (Fig. S3). Similar results were also found in a genetic knockout of GBA1, where GBA2 inhibition decreases endolysosomal pH suggesting that very low or no residual GBA1 can lead to storage. Homozygous GBA1 mutants where active GCase does not exist in the ER or cytosol, GBA2 inhibition corrects endolysosomal pH and size (Fig. 1B, S3).

The proposed model has a variety of therapeutic implications. The hypothesis favours inhibitory chaperones as these inhibit any mislocalised GCase activity. However, the hypothesis does not preclude the successful reversal of GBA-PD with allosteric GCase chaperones if they successfully re-direct residual enzymes away from the ER/cytoplasm. However, allosteric GCase chaperones would be expected to be less efficient and only effective over longer timescales. The hypothesis would also suggest that GlcCer synthesis is unlikely to be a successful strategy in Parkinson’s disease as further decreases in cytosolic glycolipids could potentially lead to exacerbation of defective vesicle pH (16,17).

The current hypothesis does not prohibit other gain of function mechanisms mainly relying on inappropriate protein-protein interactions (47–49). Recently it has been reported that the fully mature lysosomal form of GBA1, at least when overexpressed, associates with a variety of mitochondrial proteins (48). Since ceramide and GlcCer have been localised to the ER, GlcCer depletion affecting mitochondria should be ruled out where GCase is overexpressed. The current results may also be consistent with the changes in complex glycolipids found in Parkinson’s tissues as changes in cytoplasmic GlcCer may be related to tissue-specific and distinct synthetic pathways *via* non-vesicular GlcCer transport (4,29). It should be noted that previous interpretations of altered higher glycolipid levels in different cell lines was coincidental and where due to ‘glycolipid drift’ between different individuals (4). It is still an open question how mislocalised GCase activity leads to dysfunction. Possibilities include decreased cytoplasmic GlcCer, uncontrolled generation of ceramide or the production of GlcChol (50) which is a minor product of GCase activity. Any of these mechanisms could lead to the disruption of ER/cytosol-facing lipid rafts and are reliant on adequate levels of GlcCer (3).

## Experimental Procedures

### Materials

The reagents used in this study were obtained from Thermo Fisher unless otherwise stated. Cell media and supplements were from Gibco. LysoSensor yellow/blue BODIPY-LacCer and LysoTracker red were obtained from Invitrogen. U18666A was from Affinity Research. Chemicals (Exeter, UK). AMP-DNJ was prepared as previously described (51). Conduritol B-epoxide (CBE) were from Cayman chemicals. adaGlcCer and adaGalCer were from Matreya LLC and a kind gift from Prof Cliff Lingwood, University of Toronto.

### Cell culture and treatments

Detailed information about different cell lines used in this study mentioned in Table 1. The cells were cultured in DMEM (fibroblasts), RPMI (RAW) and IMDM (HAP1S) in 10% FCS containing 50 U/ml penicillin/streptomycin. U18666A was used at a final concentration of 300 nM, dissolved in ethanol at 1000X concentration. AMP-DNJ was added at 1-10nM and was dissolved in DMSO at 500X concentration. CBE was used at 25µM dissolved in ethanol at 1000x concentration. adaGlcCer and adaGalCer were prepared in DMSO at 40mM. U18666A and CBE were dissolved in etahonol and used at a final concentration of 300 nM and 25 µM, respectevetly. DMSO was used to dissolve AMP-DNJ, adaGlcCer and adaGalCer which were used at a final concentration of 10nM and 40 µM, respectively.

**Table 1.**
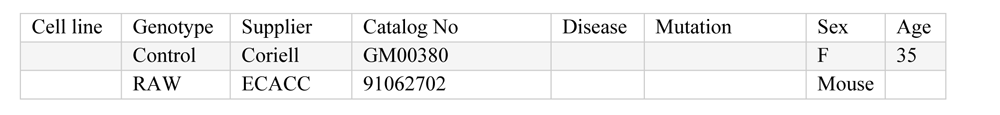
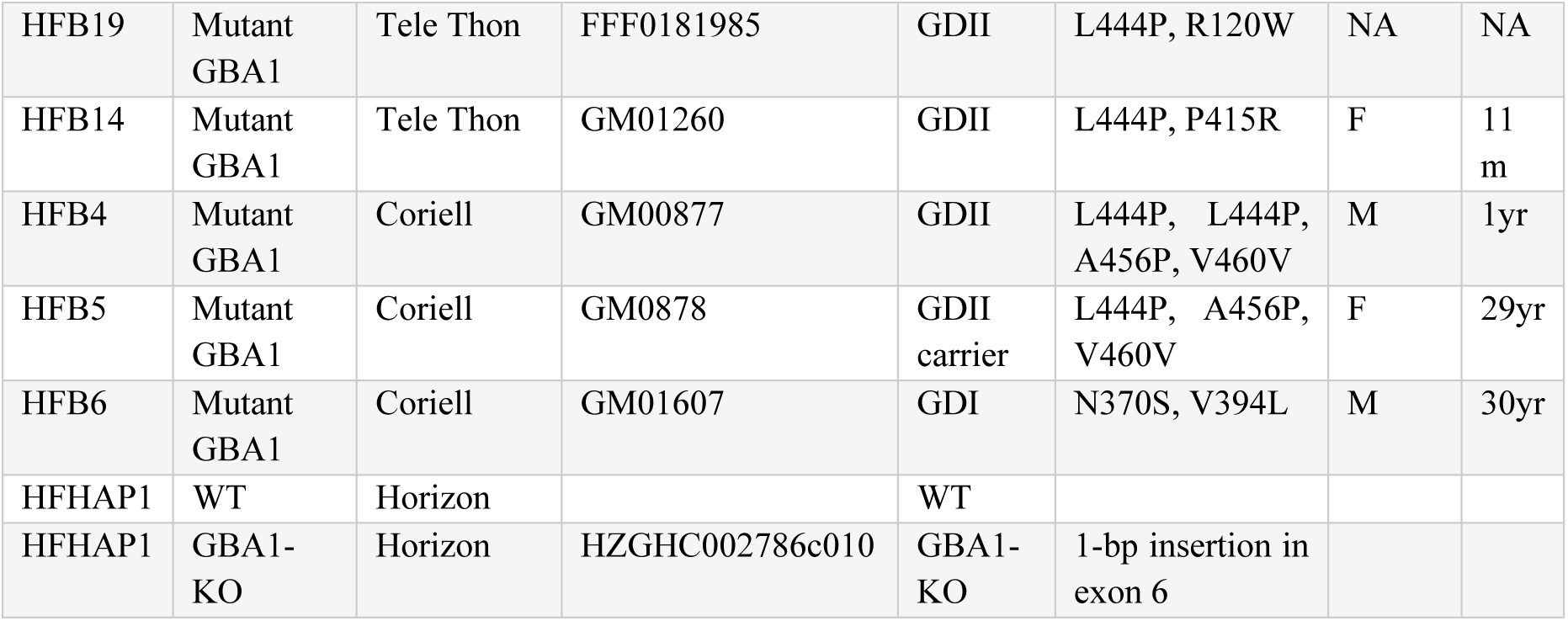
cell lines-.

| Cell line | Genotype | Supplier | Catalog No | Disease | Mutation | Sex | Age |
| --- | --- | --- | --- | --- | --- | --- | --- |
|  | Control | Coriell | GM00380 |  |  | F | 35 |
|  | RAW | ECACC | 91062702 |  |  | Mouse |  |
| HFB19 | Mutant GBA1 | Tele Thon | FFF0181985 | GDII | L444P, R120W | NA | NA |
| HFB14 | Mutant GBA1 | Tele Thon | GM01260 | GDII | L444P, P415R | F | 11 m |
| HFB4 | Mutant GBA1 | Coriell | GM00877 | GDII | L444P, L444P, A456P, V460V | M | 1yr |
| HFB5 | Mutant GBA1 | Coriell | GM0878 | GDII carrier | L444P, A456P, V460V | F | 29yr |
| HFB6 | Mutant GBA1 | Coriell | GM01607 | GDI | N370S, V394L | M | 30yr |
| HFHAP1 | WT | Horizon |  | WT |  |  |  |
| HFHAP1 | GBA1-KO | Horizon | HZGHC002786c010 | GBA1-KO | 1-bp insertion in exon 6 |  |  |

### Endolysosomal pH and size measurement

Endolysosomal pH was measured with LysoSensor yellow/blue DND-160 solution (Invitrogen, L7545, 1mM in DMSO) according to previously published method (52). A detailed protocol is available on request but briefly equal numbers of cells were used for measurement and pH curves. pH curves were generated with monensin and nigericin which were found to work variably at 0C and so were generated on a daily basis. Fluorescence data was taken at multiple times up to 40 min until stable data was obtained.

Endolysosomal size was determined using LysoTracker red (Invitrogen L7528, 1mM in DMSO). Fibroblasts were incubated with 150nM LysoTracker red (15 minutes) and washed 3 times with media. To quantify lysotracker red pixels, images were acquired by a fluorescence microscope and red pixels per cell were calculated using ImageJ.

### Lipid analysis

The lipid analysis was conducted in healthy and Gaucher patient fibroblasts as previously published (36) and protocol is available at https://www.protocols.io/view/analysis-of-glycosphingolipids-from-human-plasma-busvnwe6.

### Statistical analysis

Results are presented as a mean ±SD. In some cases, the data were normalised to the control values. Student’s one-or two-tailed t-tests and two-way ANOVA were calculated using Microsoft Excel and GraphPad prism. Significance was considered for *P*<0.05.

## Acknowledgment

We are grateful to Schlumberger foundation faculty (MB) and DMU PhD studentship (ZK).

## Additional information

**Table S1.** Glycosphingolipid levels in GBA1 mutant and healthy control with treatments, n=3.

| (nmol/ml) | GD (L444P/P415R) |  |  | HF |  |  |
| --- | --- | --- | --- | --- | --- | --- |
|  | Untreated | 10nM AMP-DNJ | 25uM CBE | Untreated | 10nM AMP-DNJ | 25uM CBE |
| GlcCer | 1.01 | 1.00 | 1.5 | 0.72 | 0.5 | 0.6 |
| GlcSph | 0.002 | 0.003 | 0.02 | 0.002 | 0.001 | 0.002 |
| Sphinganine | 0.16 | 0.27 | 0.18 | 0.16 | 0.02 | 0.17 |
| Sphingosine | 0.02 | 0.05 | 0.03 | 0.02 | 0.08 | 0.03 |
| Total GSLs | 6.95 | 7.01 | 7.53 | 5.12 | 4.25 | 5.09 |
| LacCer | 0.18 | 0.17 | 0.19 | 0.09 | 0.08 | 0.09 |
| GM3 | 2.78 | 2.55 | 3.65 | 2.0 | 1.95 | 2.21 |
| GM2 | 0.57 | 0.55 | 0.79 | 0.23 | 0.20 | 0.40 |
| GM1a | 0.06 | 0.06 | 0.08 | 0.04 | 0.04 | 0.06 |
| GD3 | 0.62 | 0.84 | 0.34 | 0.48 | 0.38 | 0.17 |
| GD1a | 0.23 | 0.22 | 0.28 | 0.13 | 0.13 | 0.06 |
| GA2 | 0.04 | 0.04 | 0.09 | 0.06 | 0.006 | 0.13 |
| Gb3 | 1.67 | 1.64 | 1.48 | 1.42 | 1.05 | 1.49 |
| Gb4 | 0.42 | 0.49 | 0.32 | 0.31 | 0.20 | 0.21 |

**Table S2.** Glycosphingolipids levels in healthy control comparison to various GBA1 mutants, n=3.

| (nmol/ml) | HF | L444P/A456P/V460V | L444P/L444P/A456P/V460V | L444P/R120W | N370S/V394L |
| --- | --- | --- | --- | --- | --- |
| Sphinganine | 0.16 | 0.34 | 0.18 | 0.18 | 0.17 |
| Sphingosine | 0.02 | 0.04 | 0.03 | 0.03 | 0.02 |
| Total GSLs | 5.12 | 7.52 | 6.20 | 7.23 | 5.22 |
| LacCer | 0.09 | 0.13 | 0.13 | 0.14 | 0.09 |
| GM3 | 2.09 | 3.36 | 2.29 | 2.57 | 1.92 |
| GM2 | 0.24 | 0.76 | 0.43 | 0.49 | 0.17 |
| GM1a | 0.04 | 0.11 | 0.04 | 0.09 | 0.03 |
| GD3 | 0.49 | 0.33 | 0.99 | 0.43 | 0.44 |
| GD1a | 0.13 | 0.37 | 0.13 | 0.29 | 0.11 |
| GA2 | 0.06 | 0.11 | 0.48 | 0.00 | 0.39 |
| Gb3 | 1.42 | 1.81 | 1.30 | 2.20 | 1.72 |
| Gb4 | 0.31 | 0.27 | 0.33 | 0.35 | 0.49 |

### β-Glucosidase activity assay

The activity of cellular GBAs was analysed using 4-methylumbelliferyl-β-D-glucopyranoside (4-MUG) (sigma) as a fluorescence substrate (2.5mM) at pH 7.4. The total protein lysate was extracted using protein extraction assay stated in western Blotting protocol. 20ug of Cell lysate was transferred to a 96-well microplate and enzymatic activity were performed in duplicates. The activity was measured in Mcllvaine Buffer pH 7.4 with 0.6% deoxycholate and triton. GBAs activity was determined using 40µM adaGlcCer as an inhibitor. The reaction mixture was incubated at 37^0^C and then reaction was stopped by adding 100ul of 0.2M Glycine pH 10.5. The fluorescence was measured in plate reader at excitation 355 nm and emission 460 nm.

**Figure S1.**
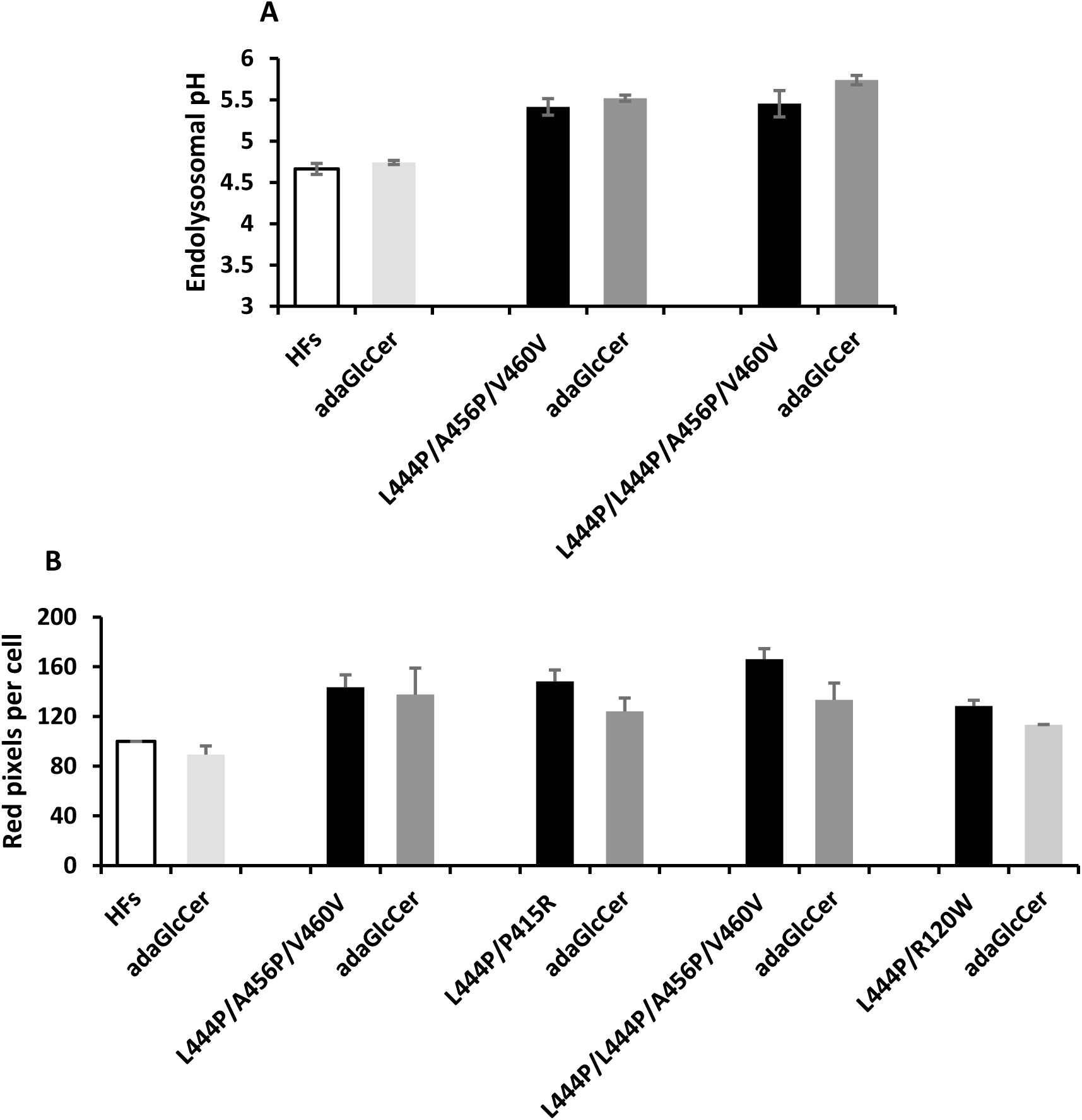
Soluble GlcCer analogue (adaGlcCer) does not correct increased endolysosomal pH and size in L444Ps GBA1 mutants. (A) Measurement of endolysosomal pH in mutant L444P GBA1 treated with 40 µM adaGlcCer. (B) quantitation of endolysosomal size in the presence of 40 µM adaGlcCer. results are presented as mean ± SD.

**Figure S2.**
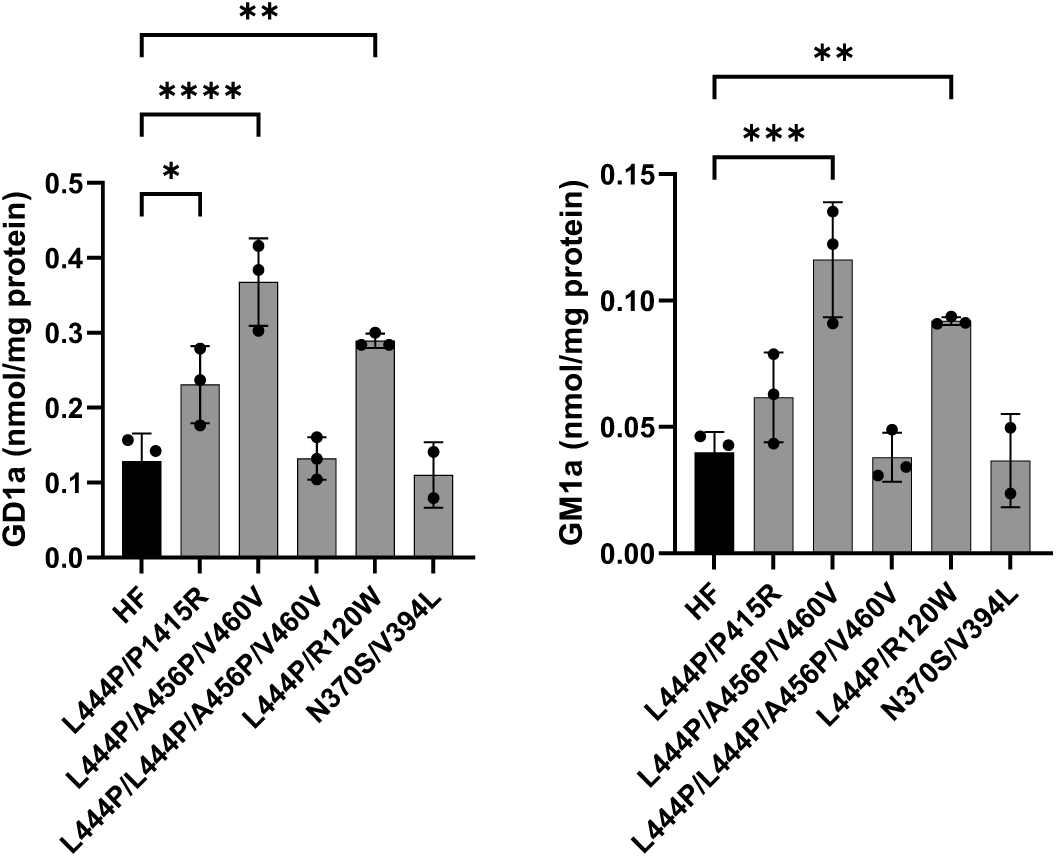
Increased a-series gangliosides in three L444P fibroblast cell lines. L444P fibroblasts show differences but not the storing homozygote. Data is represented as mean ±SD, n=3. Significance measured by t-test and ANOVA. *\*p ≤ 0.05, **p < 0.01, ***p < 0.001 and ****p < 0.001*. A lipid analysis was conducted by Danielle Taylor-Te Vruchte and Yuzhe Weng (University of Oxford).

**Figure S3.**
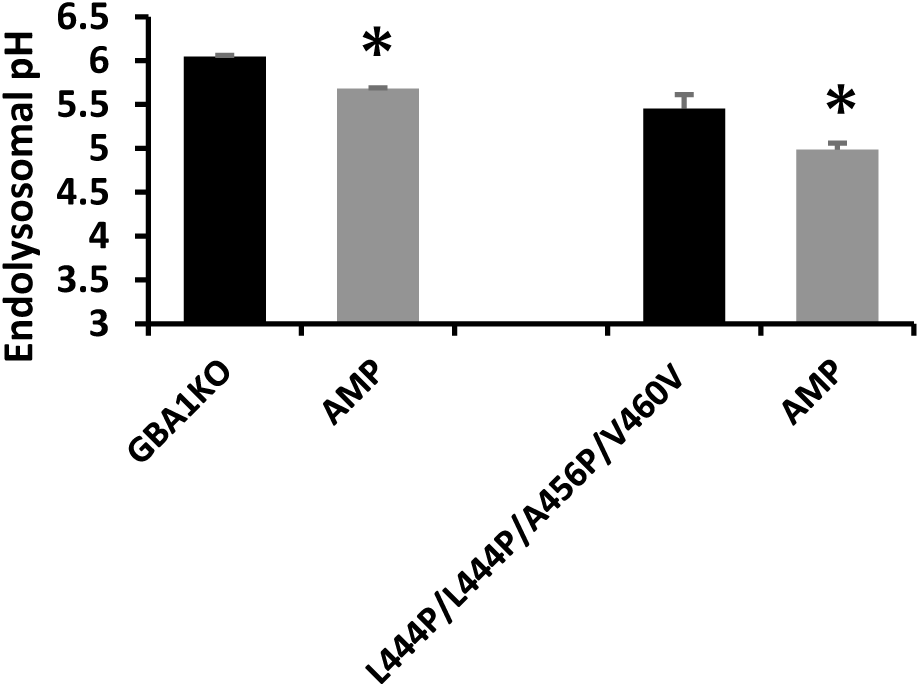
Storing Gaucher fibroblast lysosomes are sensitive to GBA2 inhibition. Severe homozygous GBA1 mutation are corrected by imino sugar inhibition of GBA2. Storing GBA1 knockouts and GBA1 L444P homozygous mutant showed decreased endolysosomal pH on inhibition of GBA2 with 10nM AMP-DNJ. All data are represented as mean ±S.D. Significance measured by t-test and ANOVA.

## References

1. Nguyen M, Wong YC, Ysselstein D, Severino A, Krainc D. Synaptic, Mitochondrial, and Lysosomal Dysfunction in Parkinson’s Disease. Trends Neurosci [Internet]. 2019;42(2):140–9. Available from: 10.1016/j.tins.2018.11.001

2. Song P, Peng W, Sauve V, Fakih R, Xie Z, Ysselstein D, et al. Parkinson&#x2019;s disease-linked parkin mutation disrupts recycling of synaptic vesicles in human dopaminergic neurons. Neuron [Internet]. 2023 Nov 3; Available from: 10.1016/j.neuron.2023.08.018

3. Smith DC, Sillence DJ, Falguières T, Jarvis RM, Johannes L, Lord JM, et al. The Association of Shiga-like Toxin with Detergent-resistant Membranes Is Modulated by Glucosylceramide and Is an Essential Requirement in the Endoplasmic Reticulum for a Cytotoxic Effect. Mol Biol Cell. 2006 Mar;17(3):1375–87.

4. Halter D, Neumann S, van Dijk SM, Wolthoorn J, de Mazière AM, Vieira O V., et al. Pre-and post-Golgi translocation of glucosylceramide in glycosphingolipid synthesis. J Cell Biol. 2007;179(1):101–15.

5. Sultana S, Stewart J, Van Der Spoel AC. Truncated mutants of beta-glucosidase 2 (GBA2) are localized in the mitochondrial matrix and cause mitochondrial fragmentation. PLoS One [Internet]. 2020;15(6):1–19. Available from: 10.1371/journal.pone.0233856

6. Wheeler S, Haberkant P, Bhardwaj M, Tongue P, Ferraz MJ, Halter D, et al. Cytosolic glucosylceramide regulates endolysosomal function in Niemann-Pick type C disease. Neurobiol Dis. 2019 Jul;127:242–52.

7. Sillence DJ. Glucosylceramide modulates endolysosomal pH in Gaucher disease. Mol Genet Metab. 2013 Jun;109(2):194–200.

8. van der Poel S, Wolthoorn J, van den Heuvel D, Egmond M, Groux-Degroote S, Neumann S, et al. Hyperacidification of Trans-Golgi Network and Endo/Lysosomes in Melanocytes by Glucosylceramide-Dependent V-ATPase Activity. Traffic. 2011 Nov;12(11):1634–47.

9. Sprong H, Degroote S, Claessens T, van Drunen J, Oorschot V, Westerink BHC, et al. Glycosphingolipids are required for sorting melanosomal proteins in the Golgi complex. J Cell Biol. 2001 Oct 29;155(3):369–79.

10. Groux-Degroote S, Van Dijk SM, Wolthoorn J, Neumann S, Theos AC, De Mazière AM, et al. Glycolipid-dependent sorting of melanosomal from lysosomal membrane oroteins by lumenal determinants. Traffic. 2008;9(6):951–63.

11. Sillence DJ, Puri V, Marks DL, Butters TD, Dwek RA, Pagano RE, et al. Glucosylceramide modulates membrane traffic along the endocytic pathway. J Lipid Res. 2002 Nov;43(11):1837–45.

12. Lesage S, Anheim M, Condroyer C, Pollak P, Durif F, Dupuits C, et al. Large-scale screening of the Gaucher’s disease-related glucocerebrosidase gene in Europeans with Parkinson’s disease. Hum Mol Genet. 2011 Jan;20(1):202–10.

13. te Vruchte D, Sturchio A, Priestman DA, Tsitsi P, Hertz E, Andréasson M, et al. Glycosphingolipid Changes in Plasma in Parkinson’s Disease Independent of Glucosylceramide Levels. Mov Disord. 2022 Oct 25;37(10):2129–34.

14. Lansbury P. The Sphingolipids Clearly Play a Role in Parkinson’s Disease, but Nature Has Made it Complicated. Mov Disord. 2022;37(10):1985–9.

15. Duran R, Mencacci NE, Angeli A V., Shoai M, Deas E, Houlden H, et al. The Glucocerobrosidase E326K Variant Predisposes to Parkinson’s Disease, But Does Not Cause Gaucher’s Disease. Mov Disord. 2013 Feb;28(2):232.

16. Schidlitzki A, Stanojlovic M, Fournier C, Käufer C, Feja M, Gericke B, et al. Double-Edged Effects of Venglustat on Behavior and Pathology in Mice Overexpressing α-Synuclein. Mov Disord. 2023;38(6):1044–55.

17. Giladi N, Alcalay RN, Cutter G, Gasser T, Gurevich T, Höglinger GU, et al. Safety and efficacy of venglustat in GBA1-associated Parkinson’s disease: an international, multicentre, double-blind, randomised, placebo-controlled, phase 2 trial. Lancet Neurol. 2023;22(8):661–71.

18. Navarro-Romero A, Fernandez-Gonzalez I, Riera J, Montpeyo M, Albert-Bayo M, Lopez-Royo T, et al. Lysosomal lipid alterations caused by glucocerebrosidase deficiency promote lysosomal dysfunction, chaperone-mediated-autophagy deficiency, and alpha-synuclein pathology. npj Park Dis 2022 81. 2022 Oct;8(1):1–15.

19. Mathieu Bourdenx, Jonathan Daniel, Emilie Genin, Federico N. Soria, Mireille Blanchard-Desce, Erwan Bezard, et al. Nanoparticles restore lysosomal acidification defects: Implications for Parkinson and other lysosomal-related diseases. Autophagy. 2016;12(3):472–83.

20. García-Sanz P, Orgaz L, Bueno-Gil G, Espadas I, Rodríguez-Traver E, Kulisevsky J, et al. N370S *-GBA1* mutation causes lysosomal cholesterol accumulation in Parkinson’s disease. Mov Disord. 2017;32(10):1409–22.

21. Zhu H, Sewell AK, Han M. Intestinal apical polarity mediates regulation of TORC1 by glucosylceramide in *C. elegans*. Genes Dev. 2015 Jun 15;29(12):1218–23.

22. Mistry PK, Liu J, Sun L, Chuang WL, Yuen T, Yang R, et al. Glucocerebrosidase 2 gene deletion rescues type 1 Gaucher disease. Proc Natl Acad Sci U S A. 2014;111(13):4934–9.

23. Marques ARA, Aten J, Ottenhoff R, van Roomen CPAA, Herrera Moro D, Claessen N, et al. Reducing GBA2 Activity Ameliorates Neuropathology in Niemann-Pick Type C Mice. PLoS One. 2015;10(8).

24. Wheeler S, Sillence DJ. Niemann–Pick type C disease: cellular pathology and pharmacotherapy. J Neurochem. 2020 Jun 15;153(6):674–92.

25. Schöndorf DC, Aureli M, McAllister FE, Hindley CJ, Mayer F, Schmid B, et al. iPSC-derived neurons from GBA1-associated Parkinson’s disease patients show autophagic defects and impaired calcium homeostasis. Nat Commun. 2014 Jun 6;5(1):4028.

26. Schonauer S, Körschen HG, Penno A, Rennhack A, Breiden B, Sandhoff K, et al. Identification of a feedback loop involving β-glucosidase 2 and its product sphingosine sheds light on the molecular mechanisms in Gaucher disease. J Biol Chem. 2017 Apr;292(15):6177–89.

27. Körschen HG, Yildiz Y, Raju DN, Schonauer S, Bönigk W, Jansen V, et al. The Non-lysosomal β-Glucosidase GBA2 Is a Non-integral Membrane-associated Protein at the Endoplasmic Reticulum (ER) and Golgi. J Biol Chem. 2013;288(5):3381–93.

28. Saito M, Rosenbergs A. The fate of glucosylceramide (glucocerebroside) in genetically impaired (lysosomal beta-glucosidase deficient) Gaucher disease diploid human fibroblasts. J Biol Chem. 1985;260(4):2295–300.

29. D’Angelo G, Uemura T, Chuang C-C, Polishchuk E, Santoro M, Ohvo-Rekilä H, et al. Vesicular and non-vesicular transport feed distinct glycosylation pathways in the Golgi. Nature. 2013;501(7465):116–20.

30. Kamani M, Mylvaganam M, Tian R, Rigat B, Binnington B, Lingwood C. Adamantyl Glycosphingolipids Provide a New Approach to the Selective Regulation of Cellular Glycosphingolipid Metabolism. J Biol Chem. 2011;286(24):21413–26.

31. Aerts JMFG, Kuo CL, Lelieveld LT, Boer DEC, van der Lienden MJC, Overkleeft HS, et al. Glycosphingolipids and lysosomal storage disorders as illustrated by gaucher disease. Curr Opin Chem Biol. 2019 Dec;53:204–15.

32. Boot RG, Verhoek M, Donker-Koopman W, Strijland A, van Marle J, Overkleeft HS, et al. Identification of the Non-lysosomal Glucosylceramidase as β-Glucosidase 2. J Biol Chem. 2007;282(2):1305–12.

33. Kuo C, Kallemeijn WW, Lelieveld LT, Mirzaian M, Zoutendijk I, Vardi A, et al. In vivo inactivation of glycosidases by conduritol B epoxide and cyclophellitol as revealed by activity-based protein profiling. FEBS J. 2019;286(3).

34. Cox TM, Schofield JP. Gaucher’s disease: clinical features and natural history. Baillieres Clin Haematol. 1997;10(4):657–89.

35. Sasagasako N, Kobayashi T, Yamaguchi Y, Shinnoh N, Goto I. Glucosylceramide and Glucosylsphingosine Metabolism in Cultured Fibroblasts Deficient in Acid β-Glucosidase Activity1. J Biochem. 1994;115(1):113–9.

36. Mylene Huebecker, Elizabeth B. Moloney, Aarnoud C. van der Spoel, David A. Priestman, Ole Isacson, Penelope J. Hallett, et al. Reduced sphingolipid hydrolase activities, substrate accumulation and ganglioside decline in Parkinson’s disease. Mol Neurodegener. 2019;14(1).

37. Jmoudiak M, Futerman AH. Gaucher disease: pathological mechanisms and modern management. Br J Haematol. 2005;129(2):178–88.

38. Ashe KM, Bangari D, Li L, Cabrera-Salazar MA, Bercury SD, Nietupski JB, et al. Iminosugar-Based inhibitors of glucosylceramide Synthase increase brain glycosphingolipids and survival in a mouse model of Sandhoff disease. PLoS One. 2011;6(6).

39. Nagatsu T, Nakashima A, Watanabe H, Ito S, Wakamatsu K, Zucca FA, et al. The role of tyrosine hydroxylase as a key player in neuromelanin synthesis and the association of neuromelanin with Parkinson’s disease. J Neural Transm [Internet]. 2023;130(5):611–25. Available from: 10.1007/s00702-023-02617-6

40. Sanchez-Martinez A, Beavan M, Gegg ME, Chau KY, Whitworth AJ, Schapira AHV. Parkinson disease-linked GBA mutation effects reversed by molecular chaperones in human cell and fly models. Sci Rep [Internet]. 2016;6(August):1–12. Available from: 10.1038/srep31380

41. Gustavsson EK, Sethi S, Gao Y, Brenton JW, García-Ruiz S, Zhang D, et al. The annotation and function of the Parkinson’s and Gaucher disease-linked gene GBA1 has been concealed by its protein-coding pseudogene GBAP1. bioRxiv [Internet]. 2023;2022.10.21.513169. Available from: 10.1101/2022.10.21.513169v2.abstract

42. Goldin E, Zheng W, Motabar O, Southall N, Choi JH, Marugan J, et al. High Throughput Screening for Small Molecule Therapy for Gaucher Disease Using Patient Tissue as the Source of Mutant Glucocerebrosidase. PLoS One. 2012;7(1).

43. Wei RR, Hughes H, Boucher S, Bird JJ, Guziewicz N, Van Patten SM, et al. X-ray and Biochemical Analysis of N370S Mutant Human Acid β-Glucosidase. J Biol Chem. 2011 Jan;286(1):299–308.

44. Steet RA, Chung S, Wustman B, Powe A, Do H, Kornfeld SA. The iminosugar isofagomine increases the activity of N370S mutant acid β-glucosidase in Gaucher fibroblasts by several mechanisms. Proc Natl Acad Sci. 2006 Sep 12;103(37):13813–8.

45. Lorent JH, Levental KR, Ganesan L, Rivera-Longsworth G, Sezgin E, Doktorova M, et al. Plasma membranes are asymmetric in lipid unsaturation, packing and protein shape. Nat Chem Biol. 2020;16(6):644–52.

46. Choy FYM. Gaucher disease: The effects of phosphatidylserine on glucocerebrosidase from normal and Gaucher fibroblasts. Hum Genet. 1984;67(4):432–6.

47. Kuo S-H, Tasset I, Cheng MM, Diaz A, Pan M-K, Lieberman OJ, et al. Mutant glucocerebrosidase impairs α-synuclein degradation by blockade of chaperone-mediated autophagy. Sci Adv. 2022 Feb;8(6):eabm6393.

48. Baden P, Perez MJ, Raji H, Bertoli F, Kalb S, Illescas M, et al. Glucocerebrosidase is imported into mitochondria and preserves complex I integrity and energy metabolism. Nat Commun. 2023;14(1):1–21.

49. Klein AD, Outeiro TF. Glucocerebrosidase mutations disrupt the lysosome and now the mitochondria. Nat Commun. 2023;14(1):10–2.

50. Akiyama H, Kobayashi S, Hirabayashi Y, Murakami-Murofushi K. Cholesterol glucosylation is catalyzed by transglucosylation reaction of β-glucosidase 1. Biochem Biophys Res Commun. 2013 Nov;441(4):838–43.

51. Overkleeft HS, Renkema GH, Neele J, Vianello P, Hung IO, Strijland A, et al. Generation of Specific Deoxynojirimycin-type Inhibitors of the Non-lysosomal Glucosylceramidase*. 1998.

52. Fraldi A, Annunziata F, Lombardi A, Kaiser H-J, Medina DL, Spampanato C, et al. Lysosomal fusion and SNARE function are impaired by cholesterol accumulation in lysosomal storage disorders. EMBO J. 2010;29(21):3607.

